# Balance training improves feedback control of perturbed balance in older adults

**DOI:** 10.1101/2021.03.31.437824

**Authors:** Leila Alizadehsaravi, Sjoerd M. Bruijn, Jaap H. van Dieën

## Abstract

Recovering balance after perturbations becomes challenging with aging, but an effective balance training could reduce such challenges. In this study, we examined the effect of balance training on feedback control after unpredictable perturbations by investigating balance performance, recovery strategy, and muscle synergies. We assessed the effect of balance training on unipedal perturbed balance in twenty older adults (>65 years) after short-term (one session) and long-term (3-weeks) training. Participants were exposed to random medial and lateral perturbations consisting of 8-degree rotations of a robot-controlled balance platform. We measured full-body 3D kinematics and activation of 9 muscles (8 stance leg muscles, one trunk muscle) during 2.5 s after the onset of perturbation. The perturbation was divided into 3 phases: phase1 from the onset to maximum rotation of the platform, phase 2 from the maximum rotation angle to the 0-degree angle and phase 3 after platform movement. Balance performance improved after long-term training as evidenced by decreased amplitudes of center of mass acceleration and rate of change of body angular momentum. The rate of change of angular momentum did not directly contribute to return of the center of mass within the base of support, but it reoriented the body to an aligned and vertical position. The improved performance coincided with altered activation of synergies depending on the direction and phase of the perturbation. We concluded that balance training improves control of perturbed balance, and reorganizes feedback responses, by changing temporal patterns of muscle activation. These effects were more pronounced after long-term than short-term training.

## Introduction

In theory, the nervous system can use two control mechanisms to recover balance after a perturbation^1^. Reactive or feedback control, occurs after a perturbation and is the only mechanism available when the nervous system has no prior knowledge of a perturbation^2,3^. Anticipatory or feedforward control is based on expectations of a perturbation, and aims to minimize the impact of the perturbation on balance by changing joint orientation or stiffness prior to a perturbation^4^. Depending on a perturbation’s direction and magnitude, feedforward control is not always sufficient for balance control, and then feedback control comes into play to regain balance. Effective feedforward control minimizes the effect of perturbations and reduces the need for feedback control^5,6^.

Three movement strategies are well known to contribute to feedback control of balance after perturbations: the ankle, counter-rotation, and stepping strategies^7^. The stepping strategy aims to displace or expand the base of support beyond the projection of the center of mass by stepping or grabbing a handhold. It is usually seen as a last resort reflecting poorer balance control, and older adults use it more than younger adults^8–10^. The ankle strategy aims to accelerate the center of mass towards the base of support through a shift of the center of pressure, the point of application of the ground reaction force, generated by ankle moments^11^. The counter-rotation strategy aims to accelerate the center of mass towards the base of support through horizontal ground reaction forces generated by changes in the angular momentum of body segments relative to the center of mass^7,11,12^. Thus, these strategies can be differentiated by distinct kinematics and kinetics but also by distinct patterns of muscle activation reflected in distinct muscle synergies^13,14^.

In non-stepping balance control, the counter-rotation strategy has been suggested to be more robust than the ankle strategy^15,16^, and the use of counter-rotation strategies relative to the ankle increases with age and the magnitude of perturbations^17–20^. Older adults rely on the counter-rotation strategy at a lower level of challenge than younger adults, even during unperturbed balancing^17,19^. This presumably helps to secure robust balance control regardless of age-related sensory errors^21,22^.

Balance training has been shown to result in altered muscle synergies and kinematics after a perturbation^23,24^. This may reflect improved feedback control but may also reflect improved feedforward control. Previously, we showed that training of older adults focusing on balance control on unstable surfaces, improved performance in perturbed and unperturbed balance tasks^25^. In addition, we found that the duration of co-contraction of muscles around the ankle increased, and we suggested that this may reflect an improved feedforward control strategy that contributed to performance improvements. Thus, training may have improved feedforward control resulting in less use of the counter-rotation strategy for balance recovery after a perturbation. However, in spite of the fact that the training program did not contain sudden, unpredictable perturbations, the challenging exercises used in training may also have improved feedback balance control, in which case one might expect the more effective counter-rotation strategy to be used more after training. In this study, we investigated the effects of training on kinematics and muscle synergies of balance recovery after perturbations in more detail to improve our understanding of training effects on feedback control of balance in older adults.

## Methods

The data collection and training were described earlier^25^, here we provide a brief summary. In this study twenty older adults (71.9±4.09 years old) participated. All participants provided written informed consent prior to participation, and the ethical review board of the Faculty of Behavioural and Movement Sciences, Vrije Universiteit Amsterdam, approved the experimental procedures (VCWE-2018-171).

Training consisted of balancing on balance boards and foam pads. The first training session was completed individually (30 minutes), and subsequently, a 3-weeks training program was completed in groups of 6-8 participants (45×3 minutes per week). We gradually increased the challenge of exercises by reducing hand support, moving from bipedal to unipedal stance, using more unstable support surfaces, and adding perturbations such as catching and throwing a ball and reducing visual input.

We assessed balance recovery with participants in unipedal stance on their dominant leg on a robot-controlled platform (HapticMaster, Motek, Amsterdam, the Netherlands). Participants performed 5 trials of a perturbed unipedal balance task, in which 12 random perturbations (6 medial and 6 lateral) were induced during 50-60 seconds. The platform rotated over a sagittal axis in the medial or lateral direction (amplitude of 8°) in random order. Participants were given two minutes rest between trials and a randomized 3-5 seconds rest period between perturbations within the trial. Participants were asked to fix their vision on a target in front of them. Full-body 3D kinematics were tracked by one Optotrak camera array (Northern Digital, Waterloo, Canada). Surface electromyography (EMG) data were recorded from nine unilateral muscles of the dominant leg: tibialis anterior (TA), vastus lateralis (VL), lateral gastrocnemius (GsL), soleus (SOL), peroneus longus (PL), rectus femoris (RF), biceps femoris (BF) and gluteus medius (GlM) and erector spinae (ES) muscles (TMSi, Twente, The Netherlands). We collected the data at baseline (Pre), after one training session (Post1), and after ten training sessions (Post2).

### Data analysis

Sixty perturbations per participant per time-point (30 medial and 30 lateral) were used to calculate all variables.

#### Perturbation Onset

The onset of the perturbations was detected through the platform’s rotation angle after synchronizing the platform, kinematics, and EMG data. Medial perturbations were defined when the platform started to rotate such that the big toe moved downward (eversion) and lateral when the big toe moved upward (inversion), and this was consistent for right- and left-leg dominant participants. A time window from 0.5 s before the onset of the perturbation to 2.5 s after the onset was selected for further analysis of all variables. For all variables 0.5 s baseline (from the start of the window until perturbation onset) was subtracted. Kinematics data were ensemble-averaged first over perturbations within a trial and then over trials per participant. The selected window was divided into three sub-windows; phase1 from perturbation onset to the maximum rotation angle of the platform, phase 2 (return to baseline) from maximum angle to 0-degree rotation angle of the platform and phase3 for 1 s after the platform returned to a 0-degree orientation.

#### Balance recovery, performance and strategy

We averaged the time series of center of mass displacement (CoM [m]), velocity (vCoM [m/s]) and acceleration (aCoM [m/s^2^]) in the frontal plane over all trials at a given time-point per subject. We calculated the positive and negative areas under the center of mass acceleration curve as an indicator of balance performance^26^. Next, we calculated total body angular momentum [kg.m^2^/s], its integral after division by the instantaneous moment of inertia to obtain a description of body orientation [degree], and the rate of change in total body angular momentum (time derivative of the total body angular momentum [kg.m^2^/s^2^]). We calculated the positive and negative areas under the curve of the rate of change of angular momentum as a second indicator of performance. The positive and negative areas were estimated separately for the three phases per direction of perturbation. The counter-rotation strategy is used when the rate of change in angular momentum accelerates the center of mass towards the base of support. Independent of this, angular momentum may be changed to regain upright body orientation. To assess how angular momentum changes were used, we compared the direction and timing of changes in the rate of change of angular momentum with CoM position and with body orientation.

#### Muscle synergies

EMG data were high-pass (35 Hz, bidirectional, 3rd order Butterworth) and notch filtered (50 Hz and its harmonics up to the Nyquist frequency, 1 Hz bandwidth, bidirectional, 1st order Butterworth). The filtered data were Hilbert transformed, rectified, and low-pass filtered (20 Hz, bidirectional, 3rd order Butterworth). Each rectified EMG signal was normalized to the maximum EMG value obtained over five perturbation trials per participant per time-point. The EMG data were down sampled to the sampling rate of the balance platform (100 Hz) to speed up the calculations. Subsequently, windows were selected from 0.5 s before until 2.5 s after the onset of perturbation. All windows for all participants and time-points were concatenated per perturbation direction. From these concatenated data, synergies were decomposed into a weighting matrix (spatial component) and activation profiles (temporal component) using non-negative matrix factorization (NNMF). Four synergies were extracted per perturbation direction, such that reconstructed EMG data accounted for a minimum of 90% of the variance in the EMG data^27^. Subsequently, we reconstructed the temporal activation profiles using pseudo-inverse multiplication of EMG data at the original sampling rate (2000 Hz) with the spatial components resulting from the previous step, per participant per time-point. The baseline values (mean over half a second before onset) were subtracted to focus on changes in muscle activation after perturbation.

We analyzed the activation profiles by estimating the magnitude of the activation and time to the peak activation for both the positive (excitation) and negative (inhibition) parts of the curve. The magnitude of the activation profile was calculated as the positive and negative area under the time dependent activation profiles separately for three phases. Time to the peak was estimated as the time that the maximum and minimum peak of an activation profile occurred in the selected window per synergy, per direction of perturbation. Magnitudes and time to the peak activation were averaged over 30 perturbations per participant and per time-point per direction of perturbation.

## Statistics

One-way repeated-measures ANOVAs were used to identify the main effect of Time-point (Pre, Post1, Post2) on all kinematics and synergy variables per phase except for time to the peak. For time to the peak the statistical analysis was performed for the whole perturbation duration, as the peaks could have been shifted between phases after the training. Greenhouse-Geisser corrections were used when the assumption of sphericity was violated. In case of a significant effect of Time-point, post-hoc tests with Holm’s correction for multiple comparisons (Pre-Post1 and Pre-Post2) were performed. In all statistical analyses, α = 0.05 was used.

## Results

### Balance performance

In figure 1 and 2, center of mass displacement, velocity, and acceleration (left panels), as well as orientation, angular momentum, and rate of change of angular momentum (right panels) are displayed. The phases are color-coded. In phase 1, the initial change in angular momentum and center of mass acceleration are in line with the direct effects of the perturbations. However, corrective responses can be observed since acceleration and rate of change of angular momentum changed direction before the platform reached its maximum angle. In phase 2, corrective responses further counteracted the induced angular momentum. These responses did not correct the center of mass position, but rather corrected the upper body orientation to vertical. In phase 3, the platform stopped, and CoM and orientation cross the baseline and some overshoot in both occurs. In all three phases, but most obviously in phases 1 and 2, the sign of the rate of change in angular momentum would not result in accelerations that would correct CoM position as in the counter-rotation strategy, but instead corrected body orientation. This is illustrated in the drawings in figures 1 and 2. The left drawing illustrates the effect of the perturbation. The right figure illustrates the corrective response of the subject rotating the body in the opposite direction relative to the platform rotation, which induces an acceleration of the CoM in the direction of the perturbation.

**Figure 1:**
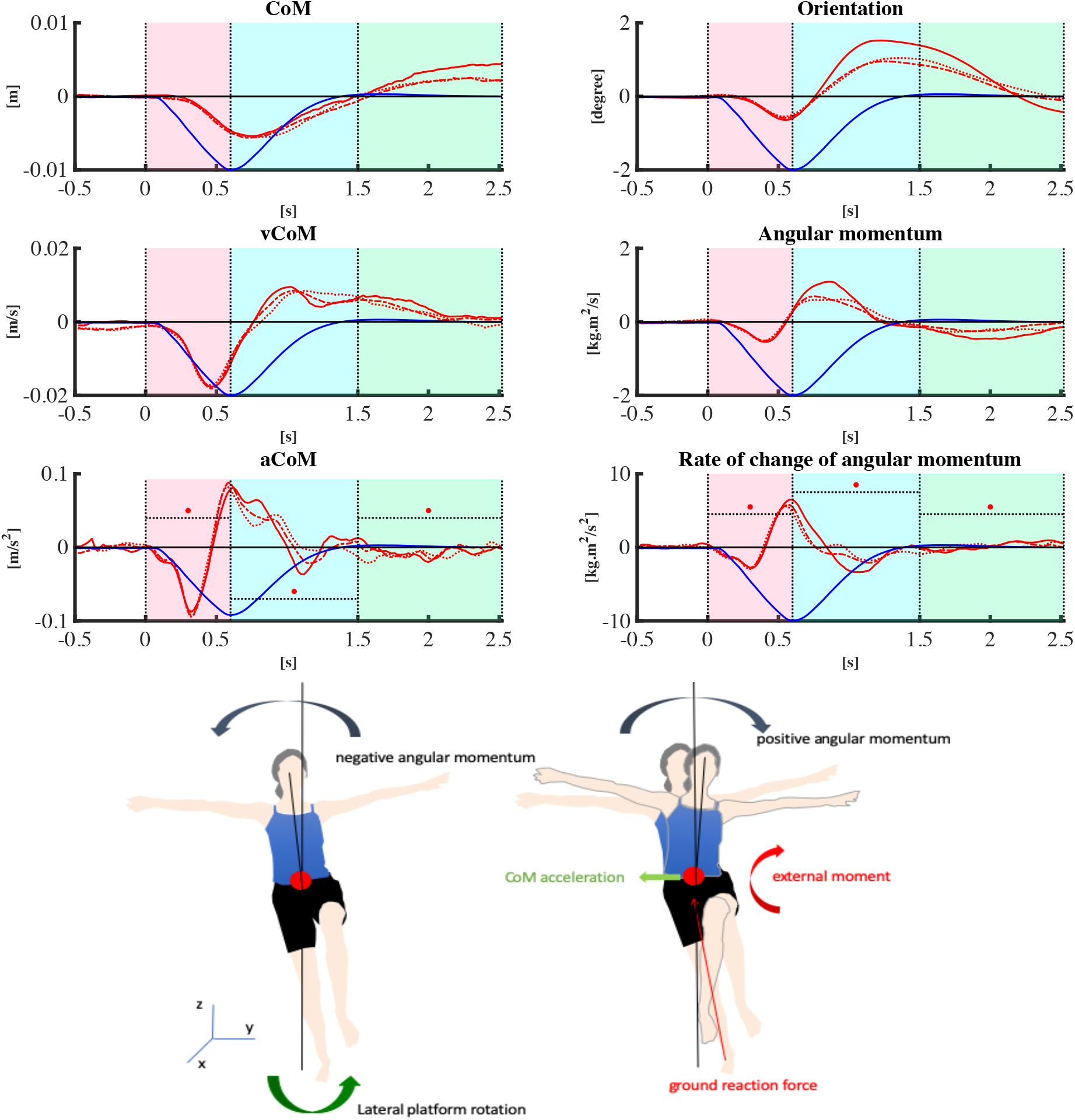
Linear kinematics (left panel) and rotational kinematics (right panel) are depicted for 0.5 s before onset to 2.5 s after onset of the lateral perturbations. Line types reflect Time-points: Pre (— solid line), Post1 (—• dash-dotted line), and Post2 (• dotted line). The red lines represent lateral perturbations; the blue line represents the rotation angle of the platform and is scaled per figure. Asterisks in 2 bottom subplots indicate a significant effect of training. The drawings illustrate the effect of the perturbation and the initial corrective response on angular momentum. The left drawing illustrates the initial, direct effect of the perturbation. The right figure illustrates the corrective response of the subject.

**Figure 2:**
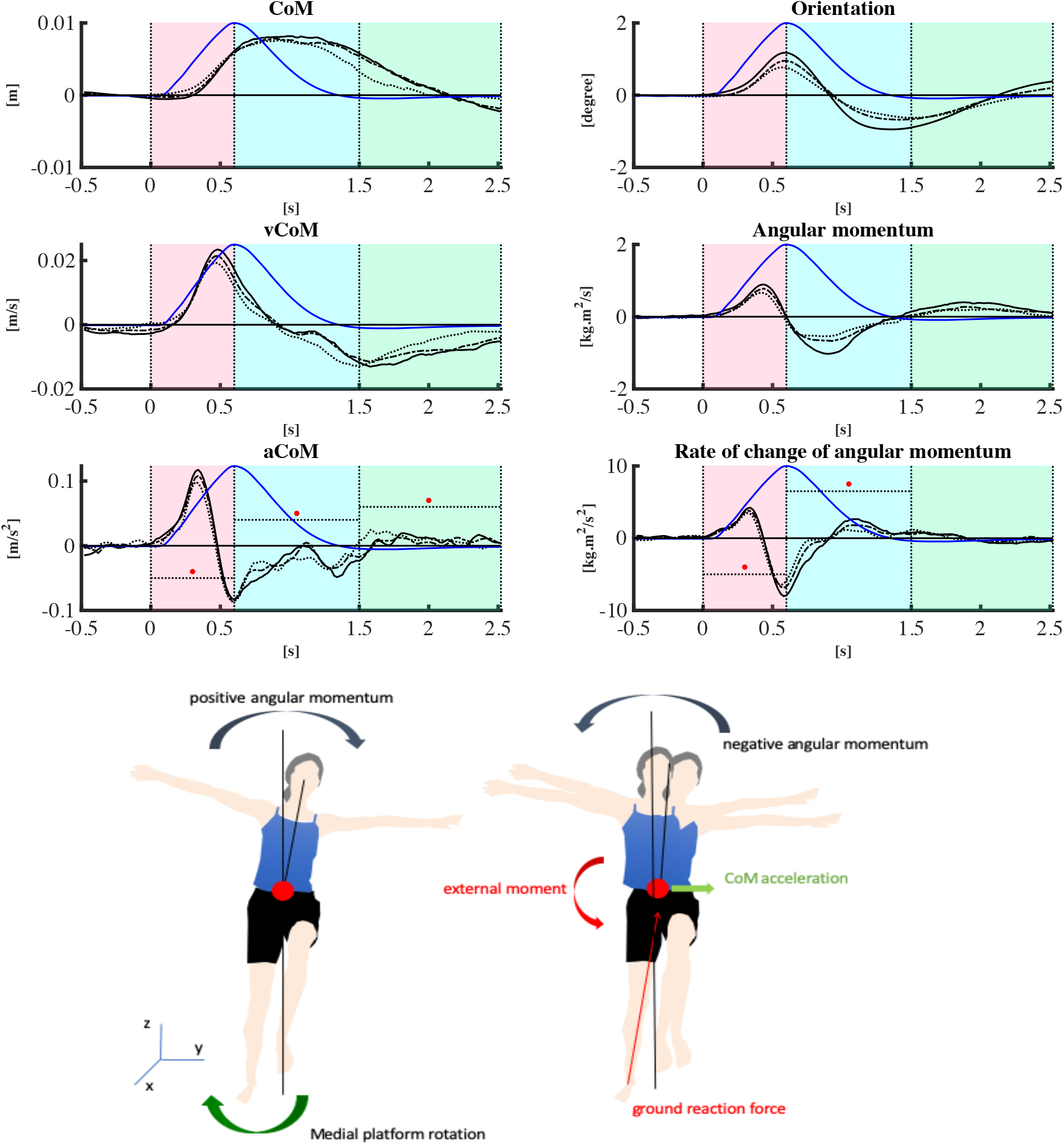
Linear kinematics (left panel) and rotational kinematics (right panel) are depicted for 0.5 s before onset to 2.5 s after onset of the medial perturbations. Lines reflect Time-points: Pre (— solid line), Post1 (—• dash-dotted line), and Post2 (• dotted line). The black lines represent the medial perturbations. The blue line represents the rotation angle of the platform and is scaled per figure. Asterisks in 2 bottom subplots indicate a significant effect of training. The drawings illustrate the effect of the perturbation and the initial corrective response on angular momentum. The left drawing illustrates the initial, direct effect of the perturbation. The right figure illustrates the corrective response of the subject.

The time series in figures 1 and 2 suggest that corrective responses were less pronounced after training, particularly after long-term training. This could be due to both improved feedforward and feedback control. Feedforward control may have affected the kinematics particularly in phase 1, where perturbation effects seemed somewhat smaller after training, especially visible after medial perturbations. In phase 2, after the training, corrective responses seemed attenuated, resulting in less overshoot in phase 3. Statistical analyses of these effect are reported below for lateral and medial perturbations separately.

#### Lateral perturbations

The negative area under the acceleration curve in phase 1, in the direction of the platform rotation, was affected by training (F_2,38 =_ 3.53, p = 0.039). Post-hoc testing showed that area under the acceleration curve did not change after short-term (p = 0.236), but decreased after long-term training (t = 2.63, p = 0.036; Figure 3.b, left panel). In phase 2, the positive area under the acceleration curve, in the direction of the platform rotation returning to horizontal, was also affected by training (F_2,38 =_ 7.46, p = 0.002). Post-hoc testing showed that area under the acceleration curve decreased after both short- and long-term training (t = 2.77, p = 0.017; t = 3.71, p = 0.002, respectively; Figure 3.a, middle panel). Also, in phase 2, the negative area under the acceleration curve in the direction opposite to the perturbation angle was affected by training (F_2,38 =_ 3.90, p = 0.029). Post-hoc testing showed that area under the acceleration curve did not change after short-term (p = 0.092) but decreased after long-term training (t = 2.66, p = 0.034; Figure 3.b, middle panel). In phase 3, the positive area under the acceleration curve was affected by training (F_2,38 =_ 9.24, p < 0.001). Post-hoc testing showed that the area under the acceleration curve decreased after both short- and long-term training (t = 3.14, p = 0.006; t = 4.11, p < 0.001, respectively; Figure 3.a, right panel).

**Figure 3:**
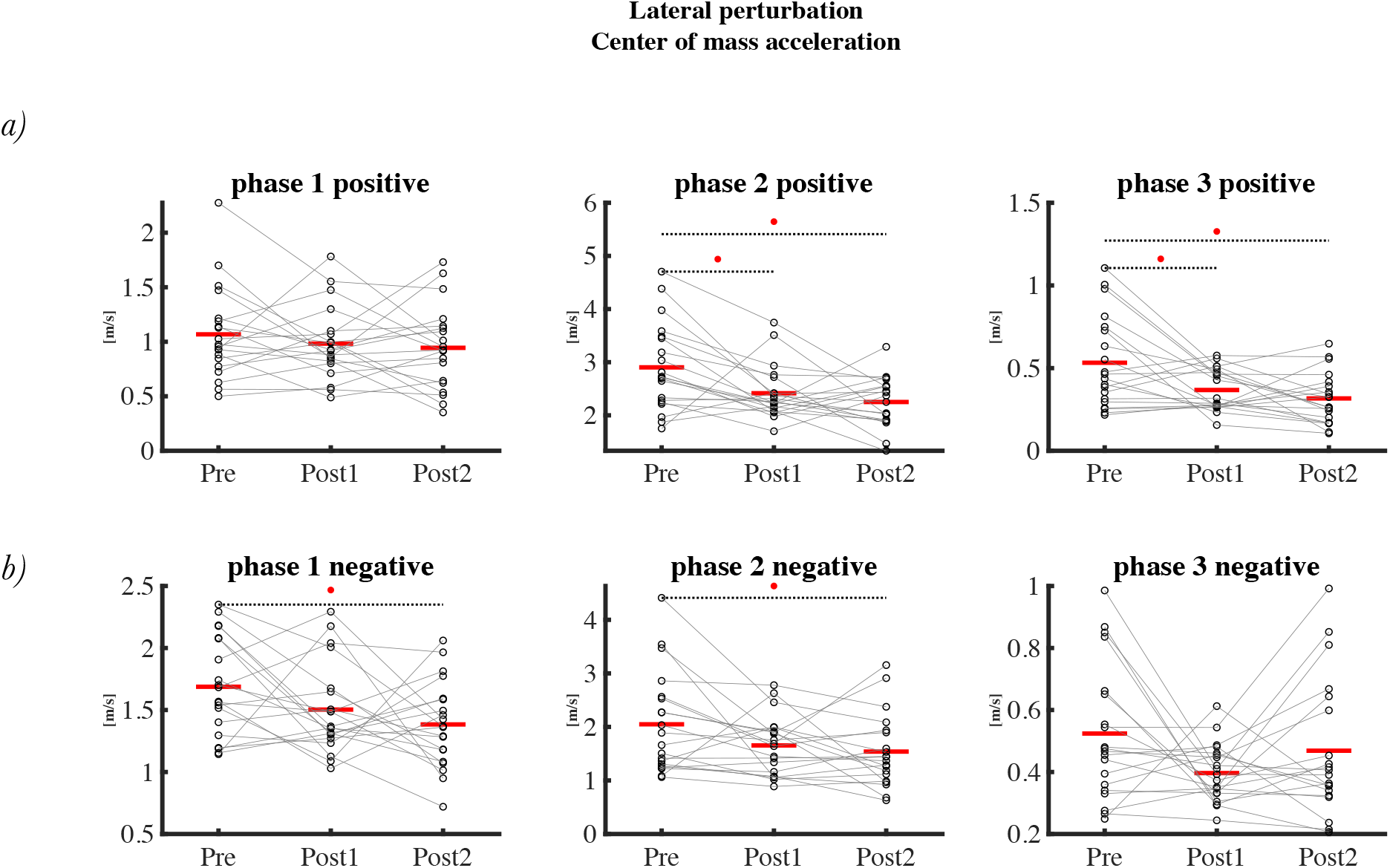
Area under center of mass acceleration curve after lateral perturbations at three time-points. Top panel a), represents the positive area, and bottom panel b) represents the negative area. Phase 1, 2 and 3 are shown in left, middle and right panel, respectively.

In phase 1, the initial negative area under the rate of change of angular momentum curve, in the direction of the platform rotation, was affected by training (F_2,38 =_ 4.52, p = 0.017). Post-hoc testing showed that area under the rate of change of angular momentum curve decreased after both short- and long-term training (t = 2.62, p = 0.038; t = 2.59, p = 0.038, respectively; Figure 4.b, left panel). In phase 2, the positive area under the rate of change of angular momentum curve, in the direction of the platform rotation returning to horizontal, was affected by training (F_1.47,28.09 =_ 11.34, p < 0.001). Post-hoc testing showed that area under the rate of change of angular momentum curve decreased after both short- and long-term training (t = 3.89, p < 0.001; t = 4.32, p < 0.001, respectively; Figure 4.a, middle panel). Also, in phase 2, the negative area under the rate of change of angular momentum curve, opposite to platform rotation, was affected by training (F_1.32,25.16 =_ 7.68, p = 0.006). Post-hoc testing showed that area under the rate of change of angular momentum curve decreased after both short- and long-term training (t = 3.27, p = 0.005; t = 3.50, p = 0.004, respectively; Figure 4.b, middle panel). In phase 3, the positive area under the rate of change of angular momentum curve was affected by training (F_2,38 =_ 4.87, p = 0.013). Post-hoc testing showed that area under the rate of change of angular momentum curve decreased after both short- and long-term training (t = 2.69, p = 0.03; t = 2.71, p = 0.03, respectively; Figure 4.a, right panel).

**Figure 4:**
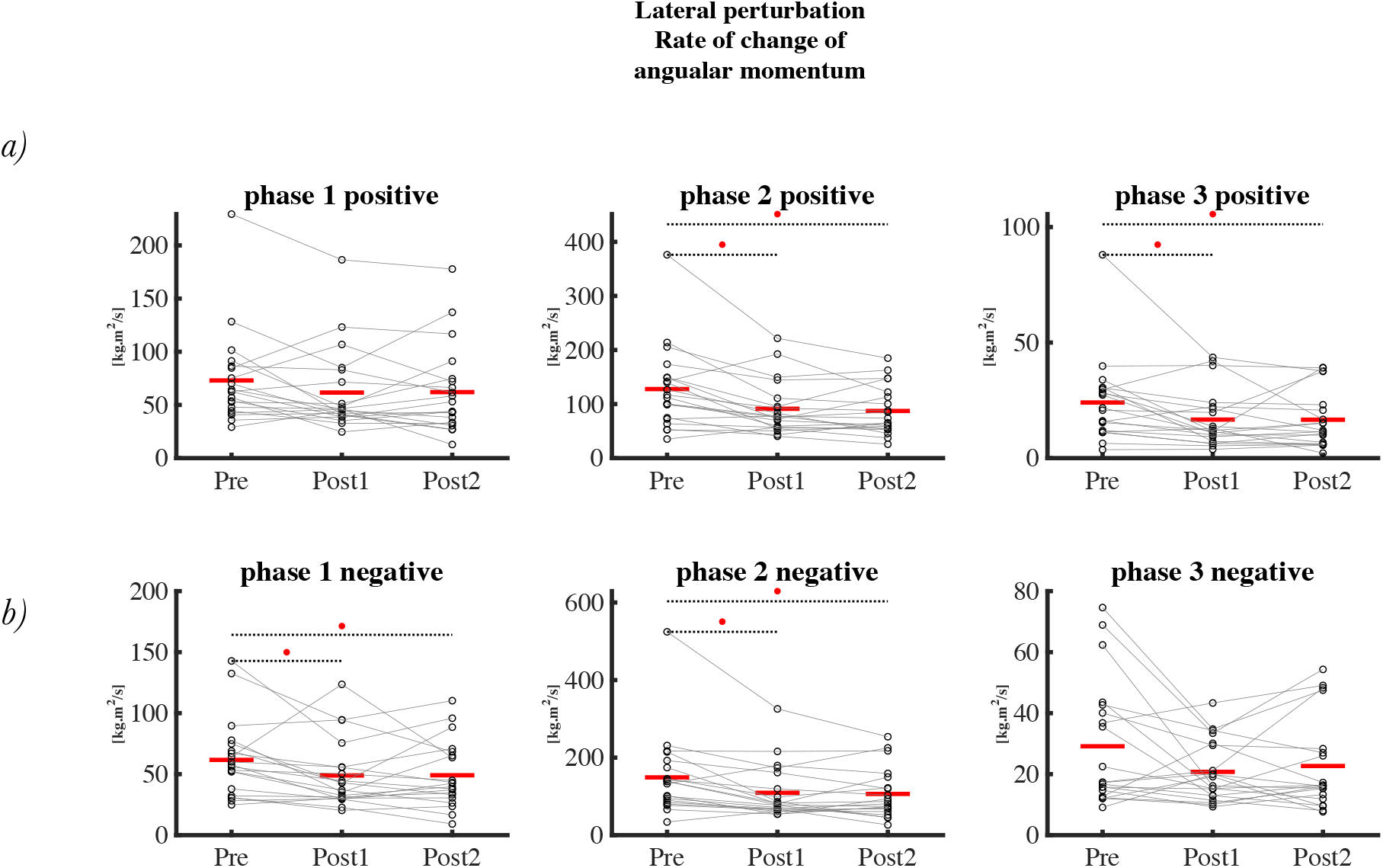
Area under the curve of the rate of change of angular momentum after lateral perturbations at three time-points. Top panel a), represents the positive area, and bottom panel b) represents the negative area. Phase 1, 2 and 3 are shown in left, middle and right panel, respectively.

#### Medial perturbations

In phase 1, the positive area under the acceleration curve, in the direction of the platform rotation, was affected by training (F_2,38 =_ 8.61, p < 0.001). Post-hoc testing showed that area under the acceleration curve did not change after short-term (p = 0.07) but decreased after long-term training (t = 4.14, p < 0.001; Figure 5.a, left panel). In phase 2, the negative area under the acceleration curve, in the direction of the platform rotation back to horizontal, was affected by training (F_2,38 =_ 7.46, p = 0.002). Post-hoc testing showed that area under the acceleration curve did not change after short-term (p = 0.092) but decreased after long-term training (t = 2.72, p = 0.029; Figure 5.b, middle panel). Also, in phase 2, the positive area under the acceleration curve, opposite to the direction of platform rotation, was also affected by training (F_2,38 =_ 3.78, p = 0.032). Post-hoc testing showed that the area under the acceleration curve did not change after short-term (p = 0.301) but decreased after long-term training (t = 2.74, p = 0.027; Figure 5.a, middle panel). In phase 3, the negative area under the acceleration curve was affected by training (F_2,38 =_ 8.63, p < 0.001). Post-hoc testing showed that area under the acceleration curve of overshoot did not change after short-term (p = 0.195) but decreased after long-term training (t = 4.072, p < 0.001; Figure 5.b, right panel).

**Figure 5:**
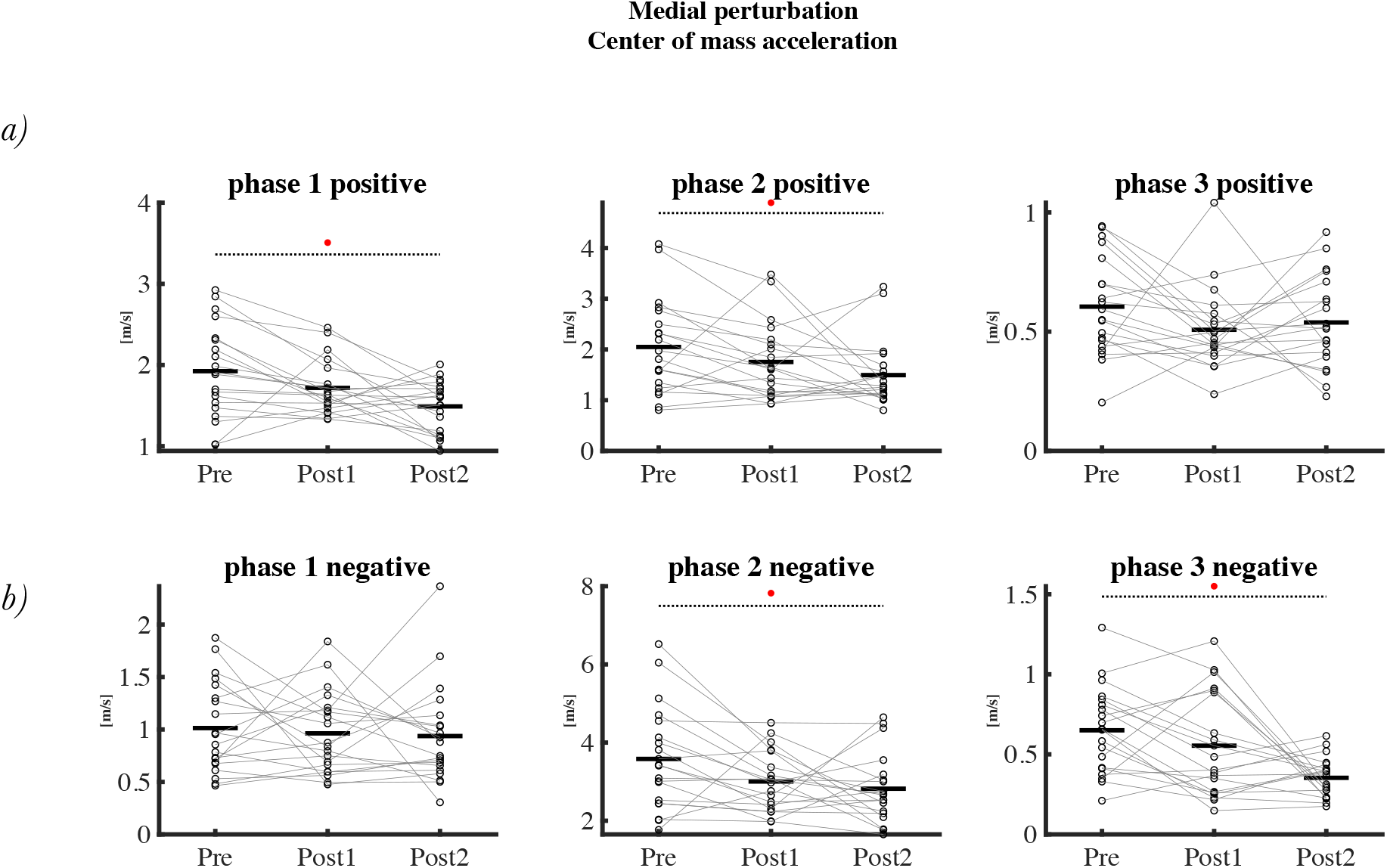
Area under the curve of the center of mass acceleration after medial perturbations at three time-point. Top panel a), represents the positive area, and bottom panel b) represents the negative area. Phase 1, 2 and 3 are shown in left, middle and right panel, respectively.

In phase 1, the initial, positive area under the rate of change of angular momentum curve, in the direction of the platform rotation, was affected by training (F_2,38 =_ 7.13, p = 0.002). Post-hoc testing showed that the area under the rate of change of angular momentum curve decreased after both short- and long-term training (t = 2.59, p = 0.027; t = 3.67, p = 0.002, respectively; Figure 6.a, middle panel). In phase 2, the negative area under the rate of change of angular momentum curve, in the direction of the platform rotation back to horizontal, was also affected by training (F_2,38 =_ 7.26, p = 0.002). Post-hoc testing showed that the area under the rate of change of angular momentum curve decreased after both short- and long-term training (t = 3.22, p = 0.005; t = 3.36, p = 0.005, respectively; Figure 6.b, middle panel). The later positive area under the rate of change of angular momentum curve in phase 2 was also affected by training (F_2,38 =_ 5.67, p = 0.007). Post-hoc testing showed that area under the rate of change of angular momentum curve decreased after both short- and long-term training (t = 3.12, p = 0.01; t = 2.65, p = 0.023, respectively; Figure 6.a, middle panel). In phase 3, no effects of training on the positive or negative area under the rate of change of angular momentum curve were found (p = 0.058, p= 0.298, respectively).

**Figure 6:**
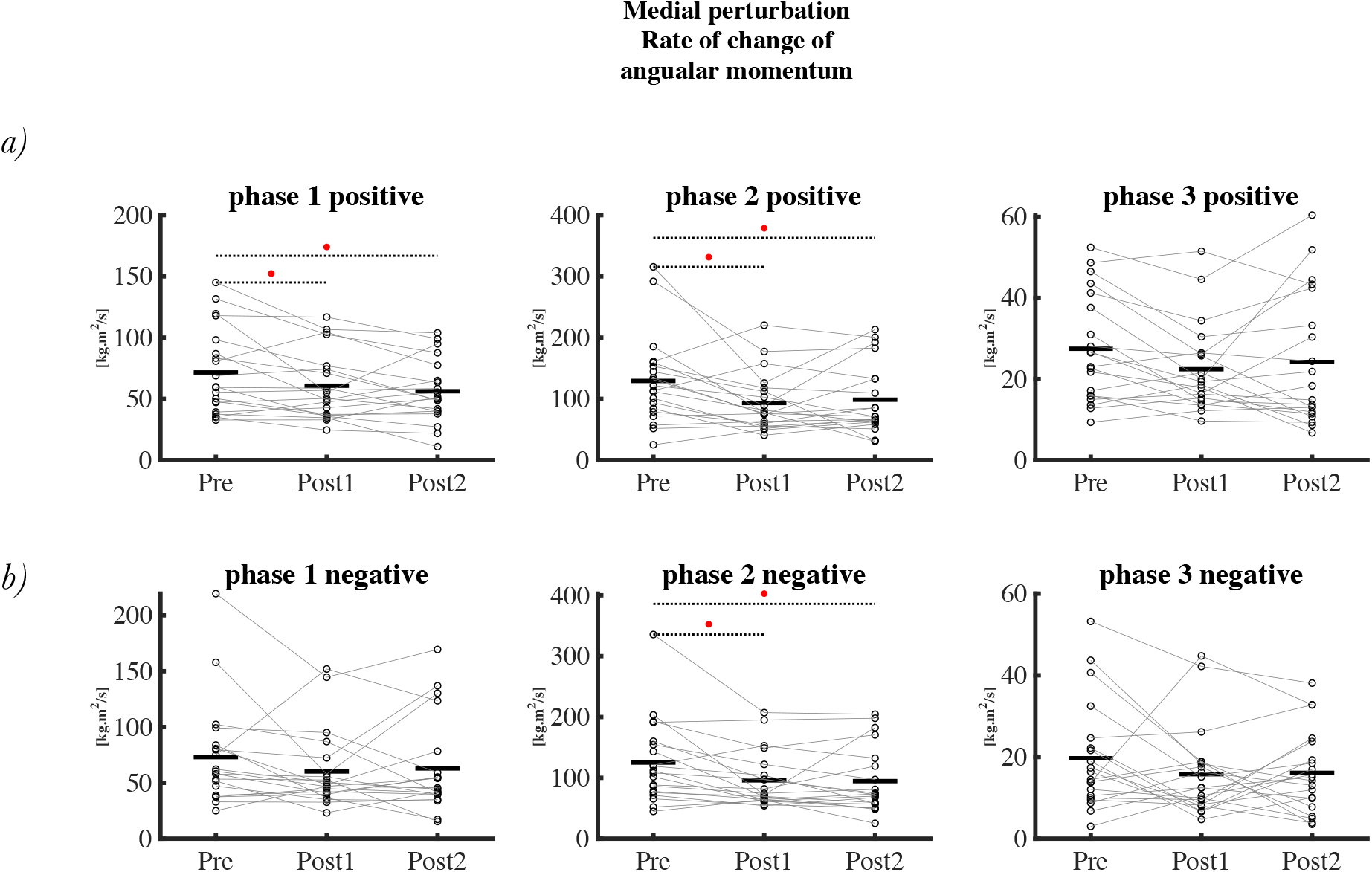
Area under the curve of the rate of change of angular momentum after medial perturbations at three time-points. Top panel a), represents the positive area, and bottom panel b) represents the negative area. Phase 1, 2 and 3 are shown in left, middle and right panel, respectively.

### Muscle Synergies

The spatial components, i.e., the weighting factors of muscles per synergy, were largely similar between medial and lateral perturbations (Figure 7). The activation profiles in synergy 2 including lateral ankle muscles and in synergy 3 including frontal ankle muscles seemed to be mirrored between lateral and medial perturbations, and therefore might reflect the use of the ankle strategy for mediolateral stabilization. But interestingly, the initial responses in these synergies, which could reflect stretch responses of the ankle muscles, would aggravate the effect of the perturbation. Synergy 1 included GlM and RF showed a fairly similar activation profile for medial and lateral perturbations and may thus be less relevant for mediolateral stabilization. Synergy 4 included the erector spinae on the stance legs side, and was mainly active after medial perturbations and may be relevant for control of upper body orientation.

**Figure 7:**
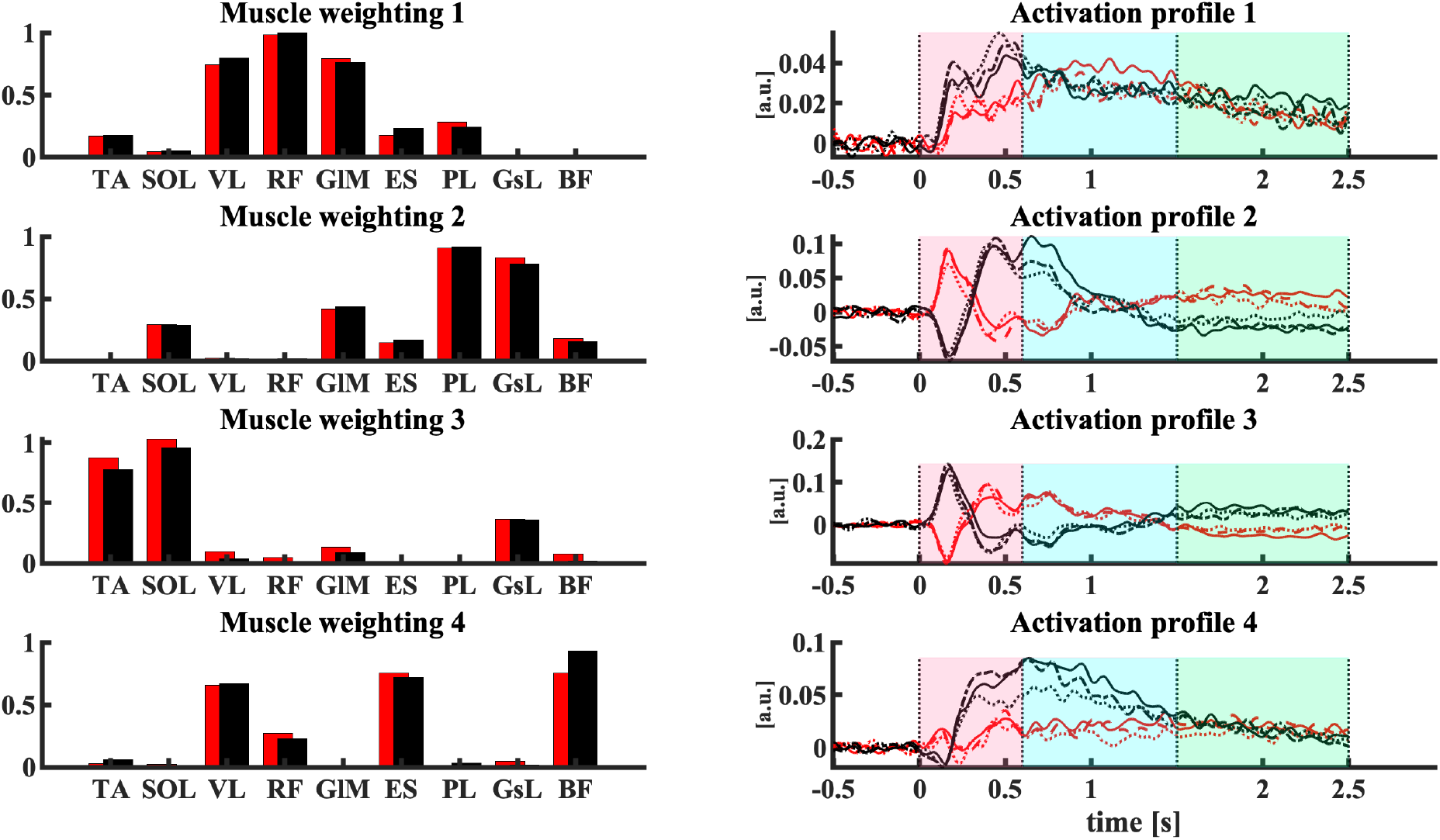
The muscle weighting and average activation profile shown after medial and lateral perturbations for three phases. Phase1 from perturbation onset to the maximum rotation angle of the platform, phase 2 (return to baseline) from maximum angle to 0-degree rotation angle of the platform and phase3 for 1 s after the platform returned to a 0-degree orientation. Lines reflect Time-points: Pre (— solid line), Post1 (—• dash-dotted line), and Post2 (• dotted line). The red color is assigned to lateral and the black color is assigned to medial perturbations. The baseline values in temporal activation in panel right were subtracted after the factorization to identify suppression and excitation relative to baseline values.

#### Lateral perturbations

In phase 1, no significant changes were observed after training.

In phase 2, although the activation profile was mainly above baseline (excitation), there was a significant effect of training on the negative area under the curve (inhibition) of synergy 1 (F_2,38 =_ 3.62, p = 0.036). However, post-hoc testing showed no significant changes after short- nor long-term training (t=2.38, p = 0.066; t=2.27, p = 0.066, respectively; Figure 8). Training also affected the negative area under the curve (inhibition) of synergy 4 in phase 2 (F_2,38 =_ 4.31, p = 0.02), although the average activation in this phase was positive. Post-hoc testing showed that activation was not changed after short-term training, but was more inhibited after long-term training (t=0.37, p = 0.70; t=2.71, p = 0.03, respectively; Figure 8).

**Figure 8:**
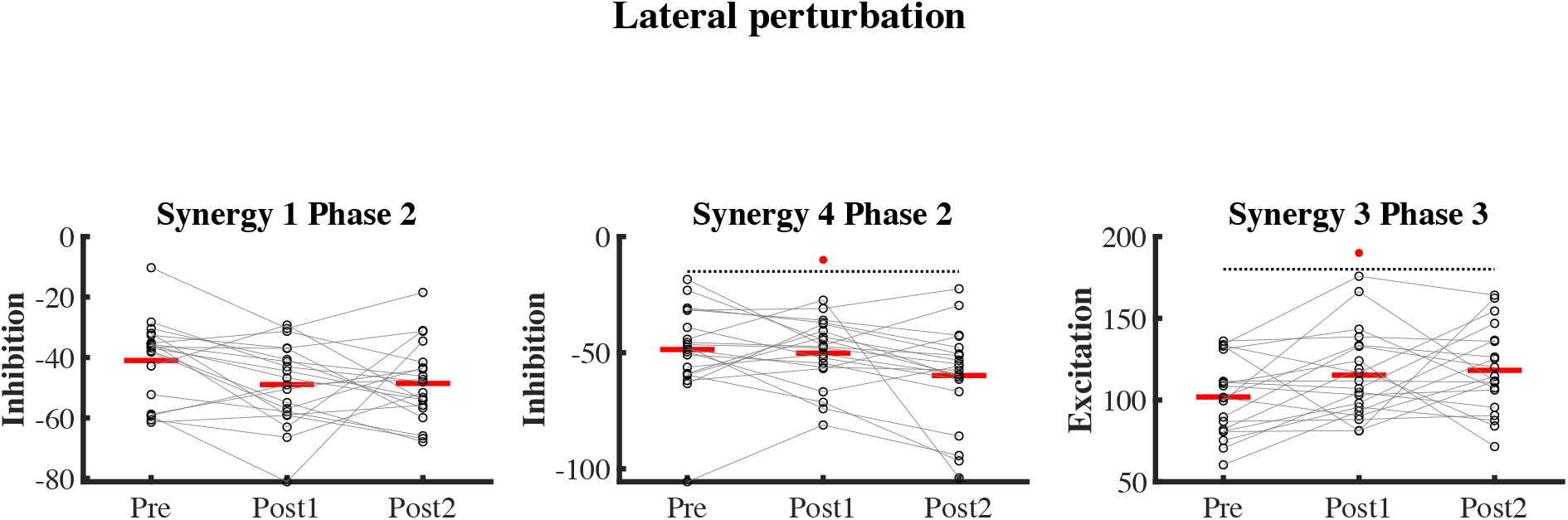
Area under the curve of activation profiles in selected phases after lateral perturbations at three time-points.

In phase 3, there was an effect of training on the positive area under the curve (excitation) of synergy 3 (F_2,38 =_ 3.67, p = 0.035). Post-hoc testing showed that the activation did not significantly change after short-term but the positive area was larger after long-term training (t = -2.08, p = 0.088; t = -2.54, p = 0.045, respectively; Figure 8).

The time to the peak did not significantly change after the training in any of the synergies.

#### Medial perturbations

In phase 1, there was an effect of training on the negative area under the curve (inhibition) of synergy 2 (F_2,38 =_ 3.52, p = 0.039) which was generally less strong after training, although post-hoc testing showed no significant differences after short-term or long-term training (t= -0.38, p = 0.70; t= -2.46, p = 0.055, respectively; Figure 9). Training increased the later negative area under the curve of synergy 3 in phase 1 (F_2,38 =_ 7.53, p = 0.002). Post-hoc testing showed that activation was more inhibited after both short- and long-term training (t=2.71, p = 0.02; t=3.76, p = 0.002, respectively; Figure 9). Training also affected the initial negative area under the curve of synergy 4 in phase 1 (F_2,38 =_ 4.99, p = 0.012). Post-hoc testing showed that activation did not change after short-term training, but was more inhibited after long-term training (t = 0.811, p = 0.422; t = 3.05, p = 0.012, respectively; Figure 9).

**Figure 9:**
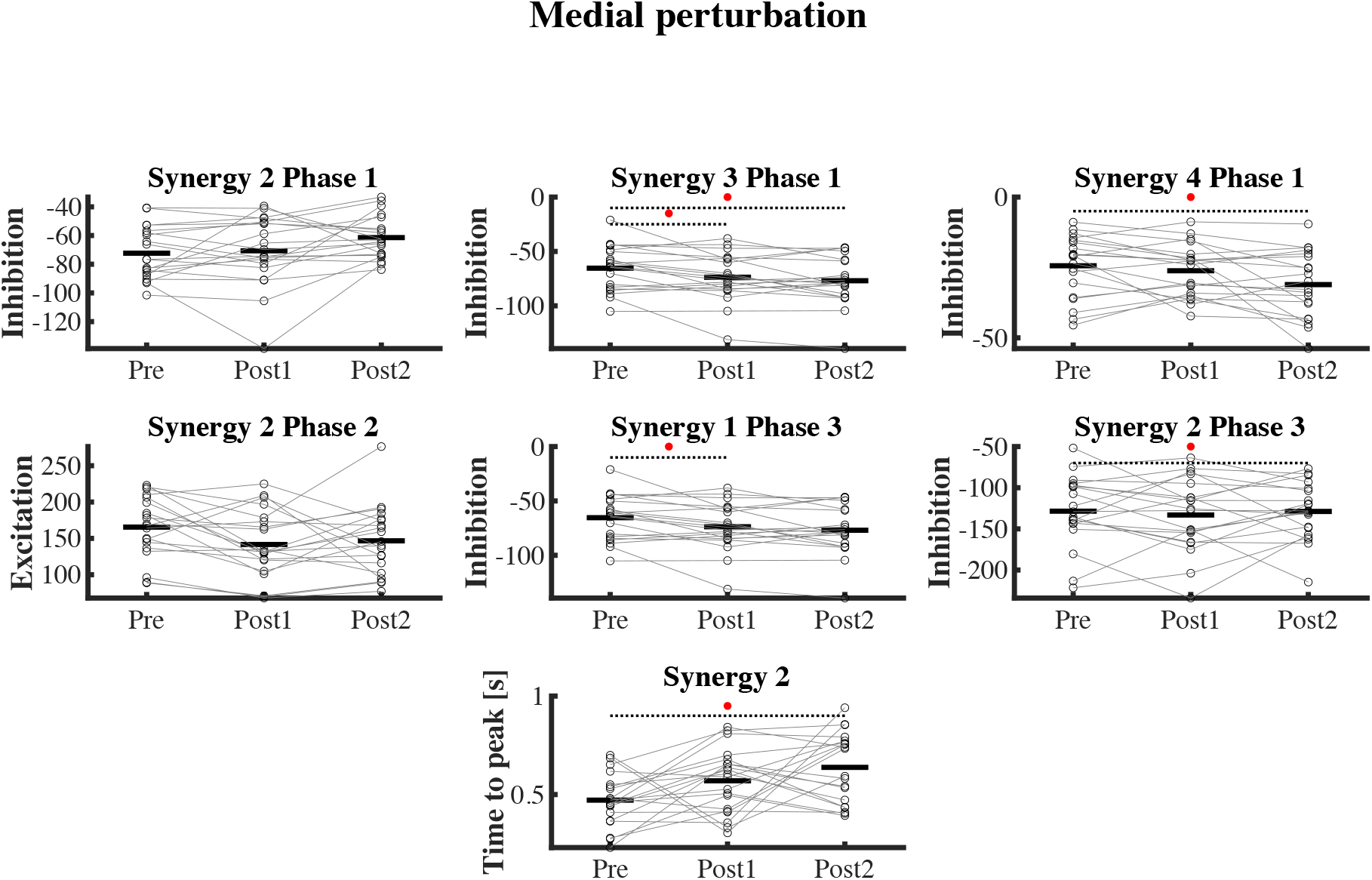
Area under the curve of activation profiles in selected phases after medial perturbations at three time-points.

In phase 2, there was an effect of training on the excitation of synergy 2 (F_2,38 =_ 3.37, p = 0.045). Post-hoc testing showed that the activation profile did not change after short-nor after long-term training (t= 2.46, p = 0.055; t=1.94, p = 0.119, respectively; Figure 9).

In phase 3, training affected the inhibition of synergy 2 in overshoot (F_2,38 =_ 3.96, p = 0.027). Post-hoc testing showed that the activation profile did not change after short-term but was less inhibited after the long-term training (t= -0.86, p = 0.39; t= -2.75, p = 0.027, respectively; Figure 9). The inhibition of synergy 1 in overshoot, phase 3, was also affected by training (F_2,38=_ 3.36, p = 0.045). Post-hoc showed it was more inhibited after short-term but back to baseline after long-term training (t = 2.53, p = 0.047; t =1.74, p = 0.177, respectively; Figure 9).

The time to the peak in synergy 2 after medial perturbation was affected by training (F_2,38 =_ 6.40, p = 0.004). Post hoc showed that time to the peak did not change after short-but was delayed after long-term training (t=-2.10, p = 0.085; t=-3.559, p = 0.003, respectively; Figure 9).

## Discussion

Aging comes with challenges to recover from perturbed balance. An effective balance training could reduce such challenges. Previously we reported that training decreased mean absolute center of mass velocity and increased ankle muscle co-contraction in perturbed unipedal balancing_25_. We suggested that increased co-contraction might compensate for the age-related deficits in sensory-motor control and as such reflect improved feedforward balance control. Yet, feedforward control is not always sufficient. Moreover, feedforward control in the form of sustained co-contraction requires energy and hence could cause fatigue. In our experiment, participants were expecting a perturbation, but were unaware of its timing and direction, allowing some, but limited feedforward control. When balance was perturbed, consistent responses in muscle activations were observed, indicating that feedback control was still used to regain balance. After training, changes in feedback control and smaller corrective responses to reorient the upper body to the upright position were observed.

Medial and lateral platform perturbations caused corresponding medial and lateral accelerations of the CoM as well as a change in angular momentum in the direction of platform rotation. Subsequent changes in angular momentum did not contribute to moving the center of mass back to the baseline position which indicates that the counter-rotation strategy was not used by our participants. Thus, the rate of change in angular momentum was used to re-orient the body rather than to shift the center of mass position over the base of support. This reorientation of the body was better tuned after training, i.e., the corrective change in angular momentum had a smaller area under the rate of change of angular momentum curve, resulting in less overshoot. This also contributed to better control of the CoM as the adverse effect on CoM acceleration would be smaller. These findings emphasize that balance control, conceptualized as control over CoM position relative to the base of support, is constrained by control of body orientation. While the CoM could be maintained over the base of support with opposite orientations of the upper and lower body oriented, a vertical orientation of both segments seems to be preferred and would of course be less demanding.

Reactive balance control improved after training, as shown by decreased amplitudes of the center of mass acceleration and rate of change of angular momentum. The improvement in center of mass acceleration and in rate of change of angular momentum in phase 1 might partially be caused by better feedforward control or improved reflex-based activity immediately following the perturbation and partially by improved feedback control in generating the corrective response at the end of the phase 1. The significant improvement in balance performance of phase 2 after perturbations indicates that the feedback control of balance improved more notably after long-term training. The improvements in phase 3 are likely an effect of better tuned responses in phase 2, resulting in less overshoot, but could also be due to higher co-contraction leading to a quicker damping of oscillations after the perturbations.

For lateral perturbations, training caused changes in the synergies in phases 2 and 3. After training and most notably after long-term training, participants inhibited synergy 4, comprising ES, VL and BF muscles, more in phase 2. This may have reduced the overshoot of CoM movement augmented by an increase in co-contraction of TA and SOL as evidenced by enhanced excitation of synergy 3 in phase 3.

For medial perturbations, training caused less inhibition in phase 1 of synergy 2 including the PL and GsL muscles. Since this initial inhibition may aggravate the perturbation, the training adaptation is likely beneficial. In the same phase the inhibition of synergy 4 including the ES, VL and BF muscles became more pronounced. The inhibition of the ES may have limited upper body rotation towards medial. Later in phase 1, synergy 3 including the TA and SOL was more inhibited and this inhibition likely helped balance recovery by the simultaneous excitation of synergy 2. In phase 2, synergy 2 was less activated after training, which may have reduced the overshoot of CoM movement, which in turn could explain the decreased inhibition of this synergy in phase 3.

## Conclusion

We investigated the effect of balance training on feedback control after expected but unpredictable balance perturbations in older adults. Our results indicate that balance training improves performance and improves the corrective responses after a perturbation. The improvement was observed in reduced amplitudes of the rate of change of angular momentum and center of mass acceleration. The rate of change of angular momentum did not correct the center of mass position, as we expected from the definition of the counter-rotation strategy, but reoriented the body to the vertical. These kinematic changes appeared to be linked to altered temporal activation of muscles grouped in ankle and upper body synergies.

## Acknowledgments

This project has received funding from the European Union’s Horizon 2020 research and innovation programme under the Marie Skłodowska-Curie grant agreement No 721577. The research team would like to thank the individuals who participated in the experiment for the purposes of this research. SMB was funded by a VIDI grant (016.Vidi.178.014) from the Dutch Organization for Scientific Research (NWO).

## Notes

### Competing Interest Statement

The authors have declared no competing interest.

